# Genotype-dependent responses to long-term water stress in *Chenopodium quinoa* Willd

**DOI:** 10.1101/2022.04.12.488037

**Authors:** I. Maestro-Gaitán, S. Granado-Rodríguez, M. Orús, J. Matías, V Cruz, L. Bolaños, M. Reguera

## Abstract

Within the current climate context, freshwater resources have become scarce. Agriculture, especially in rain-fed conditions, should deal with the need for increasing yields to contribute to food security under limiting water availability. Exploring underutilized crops such as *Chenopodium quinoa* (quinoa) has become a unique opportunity as some of these crops possess the ability to tolerate several abiotic stresses, including drought. In line with this, this work aimed at evaluating the genotype-dependent response to drought by comparing the performance of different European-adapted cultivars (F14, F15, F16, and Titicaca). The results show that the cultivars here evaluated presented different mechanisms to cope with long-term water stress, including changes in phenology, morphology, or physiology. Among them, the cultivar F16 might be the most promising genotype to grow under water-limiting conditions as it was able to increase Water Use Efficiency (WUE), reducing the stomatal conductance and keeping CO_2_ assimilation rates similar to well-watered conditions, maintaining seed yield and increasing harvest index (HI) under water deficit conditions. Furthermore, based on these results, we propose a model in which differences between a tolerant and a sensitive genotype are presented. Altogether, we believe that this work will significantly contribute to broadening our understanding regarding how quinoa responds to long-term water stress highlighting genotype-related differences that will allow the selection of the best adapted genotypes for water-limiting environments.

**Highlight:** Quinoa physiological, phenological, and morphological distinct responses to long-term water stress depending on the genotype.

## 1. Introduction

Current prospects estimate that 60% of the global population may suffer water scarcity by 2025, with drought reducing severely agriculture economic outputs (Naumann et al., 2021; Qadir et al., 2007). In line with this, in arid and semiarid areas, including those found in the Mediterranean region, water deficit is among the major constraints for agricultural production (Jacobsen et al., 2013; Tramblay et al., 2020). Hence, water limitation is threatening agriculture, with growing irrigation needs due to an increased demand for food production (Araus, 2004). Researching efficient ways to use water resources is crucial when aiming at improving water management to ensure agricultural production, securing food worldwide under changing climatic conditions (Jacobsen et al., 2013). Thus, a more efficient use of water can be achieved through the improvement of water management together with the selection of optimal crops and/or varieties for rain-fed conditions (i.e. breeding crop varieties more water-use efficient) (Araus, 2004).

*Chenopodium quinoa* Willd., commonly known as quinoa, has been widely studied in recent years due to its high nutritional value (Graf et al., 2015; Vega-Gálvez et al., 2010). It constitutes a facultative halophyte with a large genetic diversity reflecting its potential adaptability to a wide range of environments (Zou et al., 2017). In fact, it has been proposed that quinoa can be an alternative and promising crop for marginal environments as is able to tolerate well different abiotic stresses (including drought) (Choukr-Allah et al., 2016; Hinojosa et al., 2018; Jacobsen, 2003).

Singh (Singh, 2009), defined drought tolerance as the causative mechanisms of a minimum yield loss in drought conditions relative to the maximum yield obtained in an optimal environment. Thus, plants able to grow and maintain yields under limited water supplies are considered drought-tolerant (Moser, 2004). In line with this, quinoa has been defined as a drought-tolerant crop able to grow within a precipitation range that may vary between 300 and 1000 mm (with an optimal found between 500-800 mm), being (water availability) critical for the crop establishment and during seed filling stage (Gómez-Pando & Aguilar-Castellanos, 2016; Jacobsen et al., 2003). The impact of drought has been previously explored on quinoa (reviewed by Hinojosa et al. (Hinojosa et al., 2018)). In some of these aforementioned studies, the impact of severe water stress was applied at certain developmental stages, revealing that the flowering and seed filling stages are the most sensitive phases to drought and critical points determining yields in this crop (Bertero & Ruiz, 2008; Gámez et al., 2019; Hinojosa et al., 2019). Accordingly, it was shown that drought stress can accelerate quinoa flowering shortening the vegetative phase, as a mechanism to minimize dehydration, without necessarily implying yield penalties, as observed in other plant species like wheat (Jacobsen et al., 2003; Shavrukov et al., 2017). Nonetheless, differential physiological responses to drought have been observed among different quinoa genotypes in terms of yield, chlorophyll fluorescence, or CO_2_ assimilation rates supporting a genotypic role controlling water stress response in this plant species (Hinojosa et al., 2018).

Still, there are very few studies performed in quinoa analysing the physiological response to long-term water stress throughout development to assess distinct mechanisms that may be genotype-dependent. Thus, this work aimed at evaluating the physiological impact of long-term water deficit on the emergent crop quinoa throughout development, with drought stress applied from branching until seed harvesting. The experimental approach attempted to simulate western Mediterranean rain-fed conditions considering the optimal sowing date for quinoa in this particular area, which takes place in February-March, and in which the dry season (from April till the end of the life cycle) coincides with the transition to reproductive stage in this crop (Matías et al., 2021). Also, the genotypic variability linked to differential physiological responses was analysed by comparing the performance of different European-adapted cultivars.

## 2. Materials and Methods

### 2.1 Plant material, experimental design, and growth conditions

Four *Chenopodium quinoa* (quinoa) cultivars (F14, F15, F16, and Titicaca) were grown in a greenhouse located at the Centre for Plant Biotechnology and Genomics (CBGP) in Madrid, Spain (40°24’20.2”N 3°49’56.8”W). F14, F15, and F16 seeds were provided by the company Algosur S.L. (Lebrija, Spain) and the Titicaca seeds were supplied by the company Quinoa Quality (Copenhagen, Denmark).

The plants were grown under natural light conditions supplemented with high-pressure sodium (HPS) lamps from November 2020 till June 2021 (with a photoperiod varying from 9 h to 15 h light) with oscillating temperatures ranging between 15°C and 20°C. Quinoa plants were planted in 8 L pots (using a mixture peat:vermiculite (3:1) at a bulk density of 0.153 g/cm^3^ to ensure uniformity, supplemented with a controlled release fertilizer Nutricote^®^ following manufacture recommendations) and were subjected to two different water treatments: water control conditions (Well-Watered, WW), in which soil water content (SWC) was kept at 70%, and water stress conditions (Water-Deficit, WD), in which SWC was kept at 35% (Supplementary Fig. 1A) from 7^th^ week after sowing, when plants started branching.

### 2.2 Morphological parameters

Plant height was measured as the stem height, from the base part of the plant to the apical shoot. Leaf area was determined by taking images of the first pair of fully expanded leaves and then the images were processed using the open-source software ImageJ (http://rsbweb.nih.gov/ij/).

### 2.3 Plant biomass and seed yield

Plant biomass was analysed at two developmental stages, at the vegetative stage and harvesting, and was determined by cutting the plants and weighing them to measure, first, the fresh weight (FW), and then, after drying the plant material in an oven at 65°C for 72 h, to measure the dry weight (DW). Total seed yield was determined by weighting the seeds per plant at physiological maturity. Seed yield of primary panicles was separated manually to evaluate seed yield distribution along the plant. Harvest index (HI) was calculated as the ratio between the seed yield (S) and the total biomass (S + plant).

### 2.4 Photosynthetic parameters

Photosynthetic parameters were measured weekly in fully expanded leaves in the upper part of the plant. The *photosynthetic activity*, as CO_2_ assimilation rate, was determined by using a Portable Photosynthesis System (IRGA LC Pro+ ADC Bioscientific LTD, Hoddesdon, UK) at two developmental stages (pre-anthesis and at seed filling stage, that corresponded to the 13^th^ and 17^th^ week, respectively). The *chlorophyll index* was measured by using the Chlorophyll Content Meter CCM200 plus (Opti-sciences, Hudson, US). *Chlorophyll fluorescence* and *stomatal conductance* (GSW) parameters were determined by using the LI-COR Li-600 porometer and fluorometer (Lincoln, Nebraska USA). Chlorophyll fluorescence parameters were taken in light- and dark-adapted leaves (this last, after 20 minutes of dark adaptation period). The minimum chlorophyll *a* fluorescence in the dark (Fo), the maximum chlorophyll *a* fluorescence in the dark (Fm), the maximum chlorophyll *a* fluorescence in the light (Fm’), and the steady-state photosynthesis in the light (Fs) were measured and used to calculate the maximum quantum yield of photosystem II (PSII) (*Fv/Fm*), the efficiency of the PSII (*ΦPSII*), the electron transport rate (*ETR*), and non-photochemical quenching (*NPQ*). The conditions set were: a high flow rate of 150 μm.s^-1^, a match time-frequency of 10 m, a flash intensity for light-adapted leaves at 10000 μmol.m^-2^. s^-1^ and 6000 μmol.m^-2^. s^-1^ for dark-adapted leaves, a flash-length of 800 ms, leaf absorbance of 0.8, a fraction absorbance of PSII of 0.5, and an integrated modulation intensity of 6.67 μmol.m^-2^. s^-1^ for light-adapted leaves and 0.0667 μmol.m^-2^. s^-1^ for dark-adapted leaves. The integrated modulation intensity was calculated as 2*667e-9*10000 *actinic modulation rate (500 Hz for light-adapted leaves and 5 Hz for dark-adapted leaves).

### 2.5 Statistical analysis

A Three-Way ANOVA followed by a Tukey post-hoc was performed to analyse the influence of the developmental stage, water treatment, and cultivar and their interaction in the different parameters measured in this study. For variables where normality and equal variances could be assumed following a Kolmogorov-Smirnov and a Levene’s test, respectively, a One-way ANOVA test was performed, followed by a Tukey post-hoc test, to perform multiple comparisons at a probability level of 5% (*p < 0.05*). A Kruskal-Wallis test by ranks was performed when data did not present a normal distribution (tested by performing a Kolmogorov-Smirnov test (*p>0.05*)). A Welch’s ANOVA test followed by a Games-Howell post-hoc test (*p>0.05*) was performed when variances were not equal (tested by performing a Levene’s test, *p>0.05*). When data were compared by pairs, Student’s T-test or U-Mann Whitney’s test were carried out for normal or not normal data distribution, respectively. All the statistical analysis were performed using the statistical software IBM SPSS version 26.0 (IBM SPSS Inc., New York, NY, USA).

## 3. Results

### 3.1 Plant morphological responses

Differences in development appeared among cultivars and water treatments (Fig. 1). The first plants reaching the flowering stage were the Titicaca plants, independently of the water treatment applied. Under WD, Titicaca plants accelerated flowering compared to WW plants, shortening its reproductive stage (from flowering till physiological maturity) in 11 days, on average. Other differences were observed among cultivars. For instance, although WD Titicaca plants were the ones that first reached the seed filling stage, WW F14 plants were harvested three weeks earlier than the rest of WD cultivars and more than four weeks earlier than the rest of WW cultivars (Fig. 1). Also, WD F15 plants delayed their flowering 4 days, on average, compared to WW F15.

**Figure 1.**
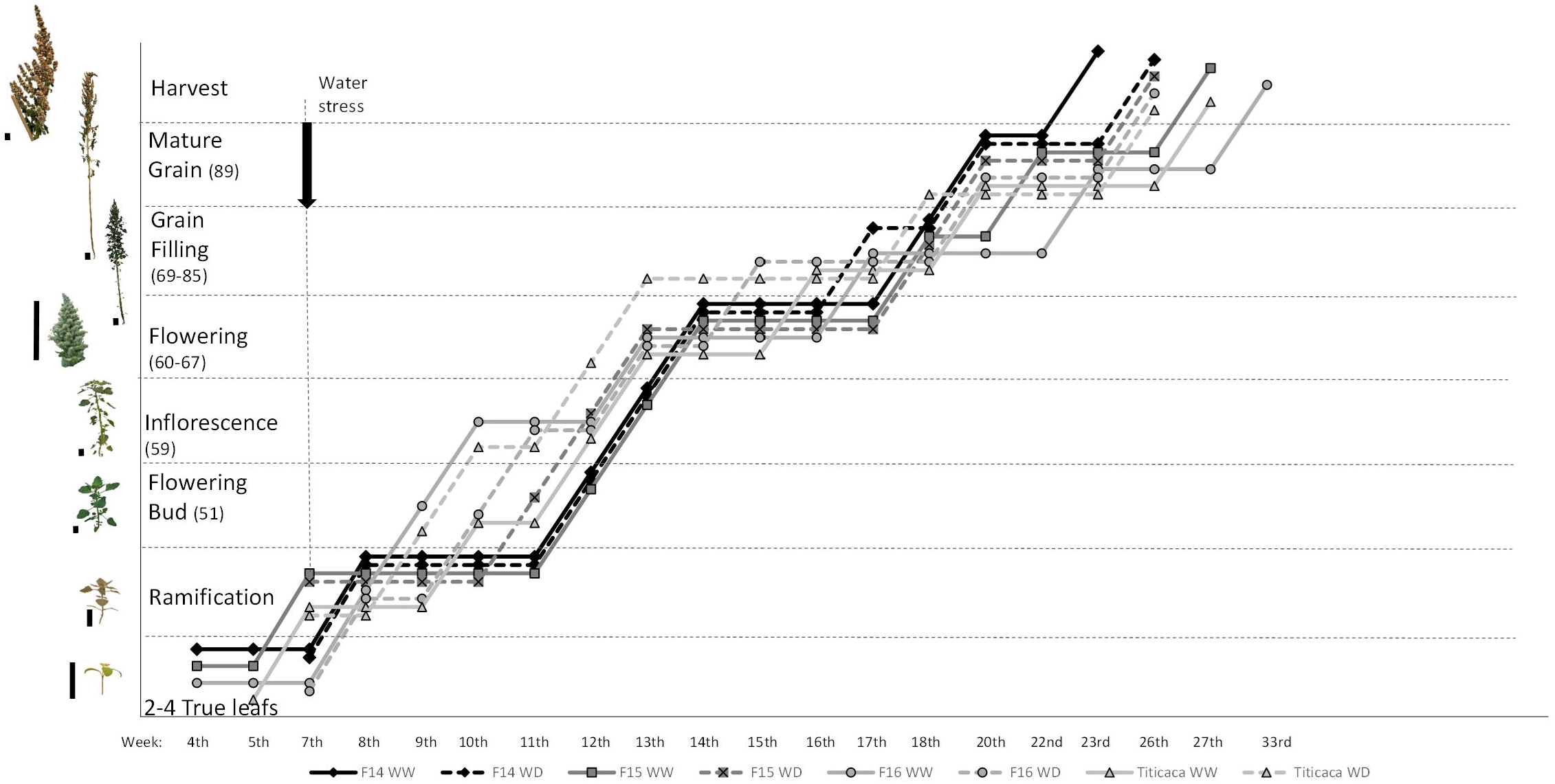
Phenological growth stages of *Chenopodium quinoa* (quinoa) cultivars grown under well-watered (WW) or water deficit (WD) conditions. Different phenological stages were identified in quinoa and the quinoa cultivars (F14, F15, F16, and Titicaca) were characterized according to the different phenological phases depending on the water treatment (WW or WD). To reach a particular phenological phase was considered when 50% or more of the plants achieved a particular developmental stage. Water stress was applied from the 7^th^ week onwards as indicated by the vertical arrow in the graph. Each water condition is represented by either continuous (WW) or dashed lines (WD) and each genotype with rhombus (F14), squares (F15), circles (F16), or triangles (Titicaca). Sample size (n) was 25. Scale bars representing the plant size are indicated as vertical lines in the left part of the images (Y axe).

The cultivar that presented the longest life cycle was F16, which lasted for 33 weeks in the case of WW plants, eight more weeks than the same cultivar growing under WD conditions. Also, F15 and Titicaca plants showed longer cycles under WW conditions contrary to F14 plants’ behaviour, presenting a longer life cycle under WD conditions.

Regarding plant height, WW plants were generally higher than those growing under WD (Fig. 2). Differences were more remarkable from week 14^th^, where plants of each condition could be grouped in two separated groups, WW and WD plants (Fig. 2A). Among genotypes, F16 plants were the tallest under both conditions (Fig 2A). These differences were maintained at harvesting (Fig. 2B), when panicle length was also measured. All cultivars presented larger panicle lengths under WW conditions compared to WD except for Titicaca, which did not show differences between treatments in this parameter (Fig. 2 C). Likewise, the cultivars that showed the largest panicles were F16 and Titicaca.

**Figure 2.**
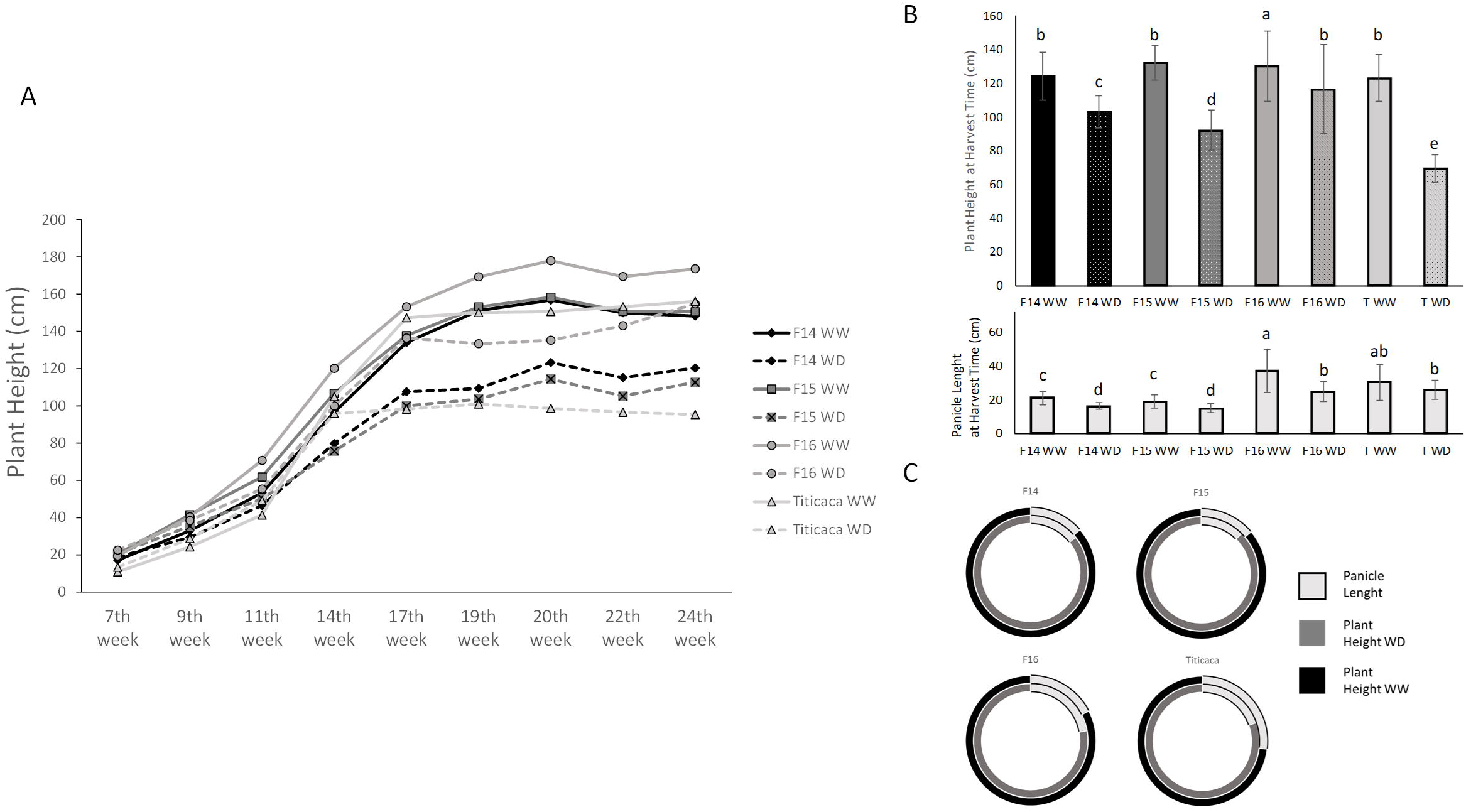
Plant height throughout development of different quinoa cultivars grown under two water regimes (WW or WD). (**A**) Plant height (cm) for each water condition represented by either continuous (WW) or dashed lines (WS) and each genotype with rhombus (F14), squares (F15), circles (F16) or triangles (Titicaca). The statistical analyses performed for this parameter are presented in Supplementary Table 2. (**B**) Plant height (upper graph) and panicle length (bottom graph) (in cm) at harvesting are presented. Columns that do not share the same letters show statistically significant differences following Kruskall-Wallis test at p-value <0.05 for both, plant height and panicle length, with n≥6. Error bars represent the standard deviation of the mean value. (**C**) The ratio between panicle length and plant height at harvesting is represented by double circles in which the external circle (black) shows the plant height under WW and the inner circle shows the plant height under WD (dark grey). Panicle length is presented proportionally to the plant height (in light grey) for each condition.

Plant biomass was first measured at the vegetative stage (at the 9^th^ week) (Fig. 3). Among the cultivars analysed grown under WW conditions, F16 and F15 plants showed larger FW than F14 or Titicaca plants, being WW F16 the one presenting the highest FW. Differences in FW appeared between water treatments in the cultivars F15 and F16, where WW plants showed higher weights (Fig. 3). On the contrary, no differences in DW appeared under WD in these cultivars, but Titicaca plants showed larger DW under WD conditions reflecting a positive impact of a water reduction in growth (Fig 3).

**Figure 3.**
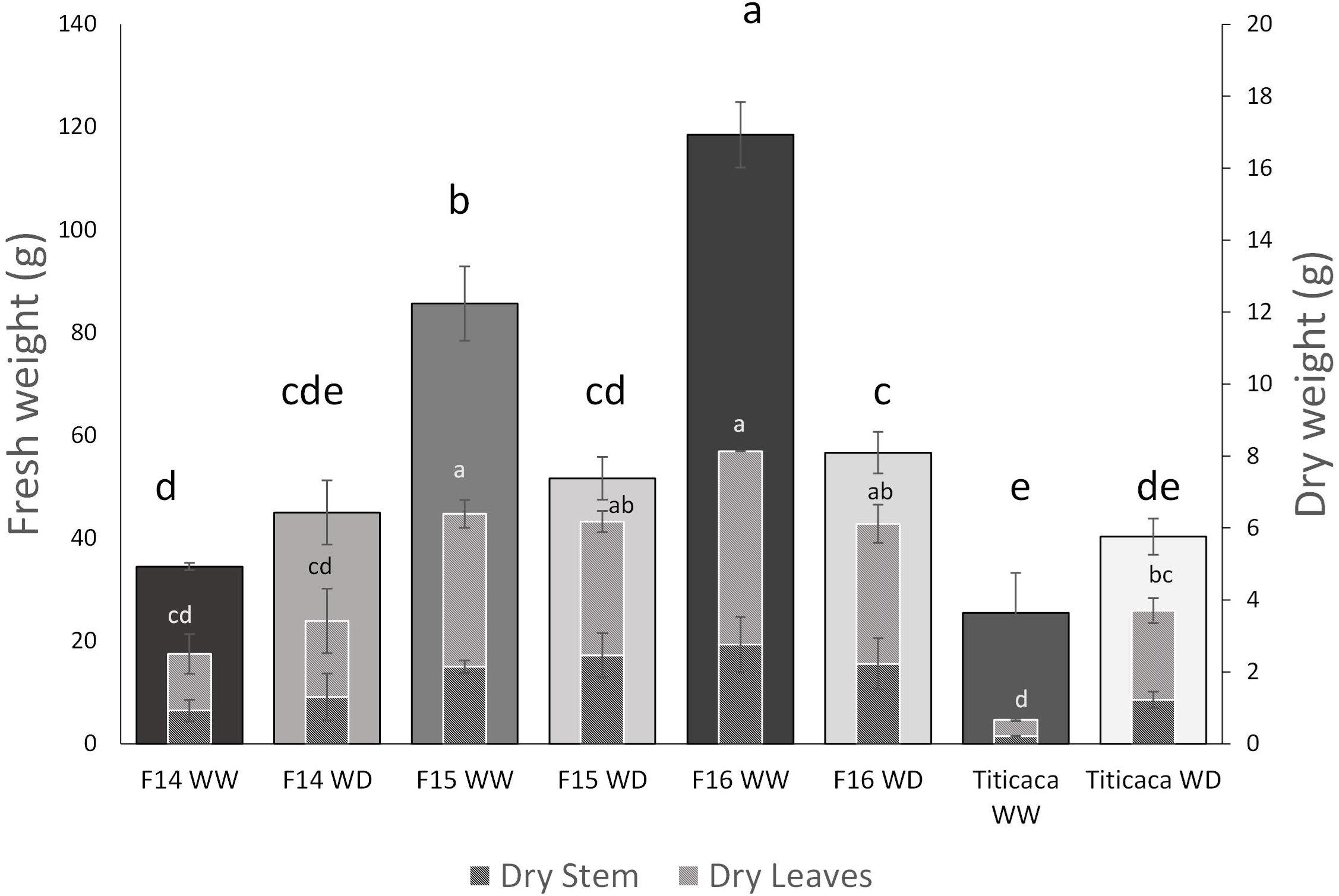
Plant biomass at vegetative stage for each quinoa cultivar and water treatment (WW or WD). Fresh weight (FW) (g) is represented by the wider columns while the dry weight (DW) (g) is represented by the inner columns, in which the stem DW (g) (dark dashed columns) and the leaves DW weight (g) (grey dashed columns) are differentiated, being the total DW of the plant the sum of both. Error bars represent the standard deviation of the mean value (n=6). Columns that do not share the same letters show statistically significant differences following Kruskall-Wallis test at *p*-value at 0.05 for FW, and One Way ANOVA followed by a Tukey post-hoc test at a *p*-value <0.05 for DW.

The ramification and number of leaves were also determined (Supplementary Table 1). F15 and Titicaca were the cultivars showing more ramifications and leaves compared to F16 or F14. It should be noted that F16 showed larger defoliation rates throughout development, under both, WW and WD, conditions. Furthermore, at the vegetative stage, all cultivars presented similar morphological characteristics, but at flowering, larger differences in the plant structure appeared. Among them, F16 plants presented a horizontal positioning of their leaves and started defoliation of bottom leaves, reaching complete defoliation of the lower half of the plant at latter stages, from seed filling stage onwards. The other cultivars presented a higher ramification number and more leaves on the lower parts of the plant, and the leaves located around the inflorescence were less turgid, showing a vertical disposition contrary to what was observed in F16 plants (Fig 4). To complement this analysis, leaf area was measured at different developmental stages (Supplementary Fig. 2). At the 2 true leaves stage, in which no water stress was yet applied, the cultivar which generated bigger fully expanded leaves was Titicaca, followed; by F15, F16, and F14. At the ramification stage (7^th^ week), no differences were found between cultivars nor water treatment. Nevertheless, when the flowering bud was emerging, differences appeared among cultivars and water treatments, being the cultivars F15 and F16 the ones showing bigger fully expanded leaves under WW conditions (Supplementary Fig. 2) and the only cultivars that reduced their leaf area under WD conditions.

**Figure 4.**
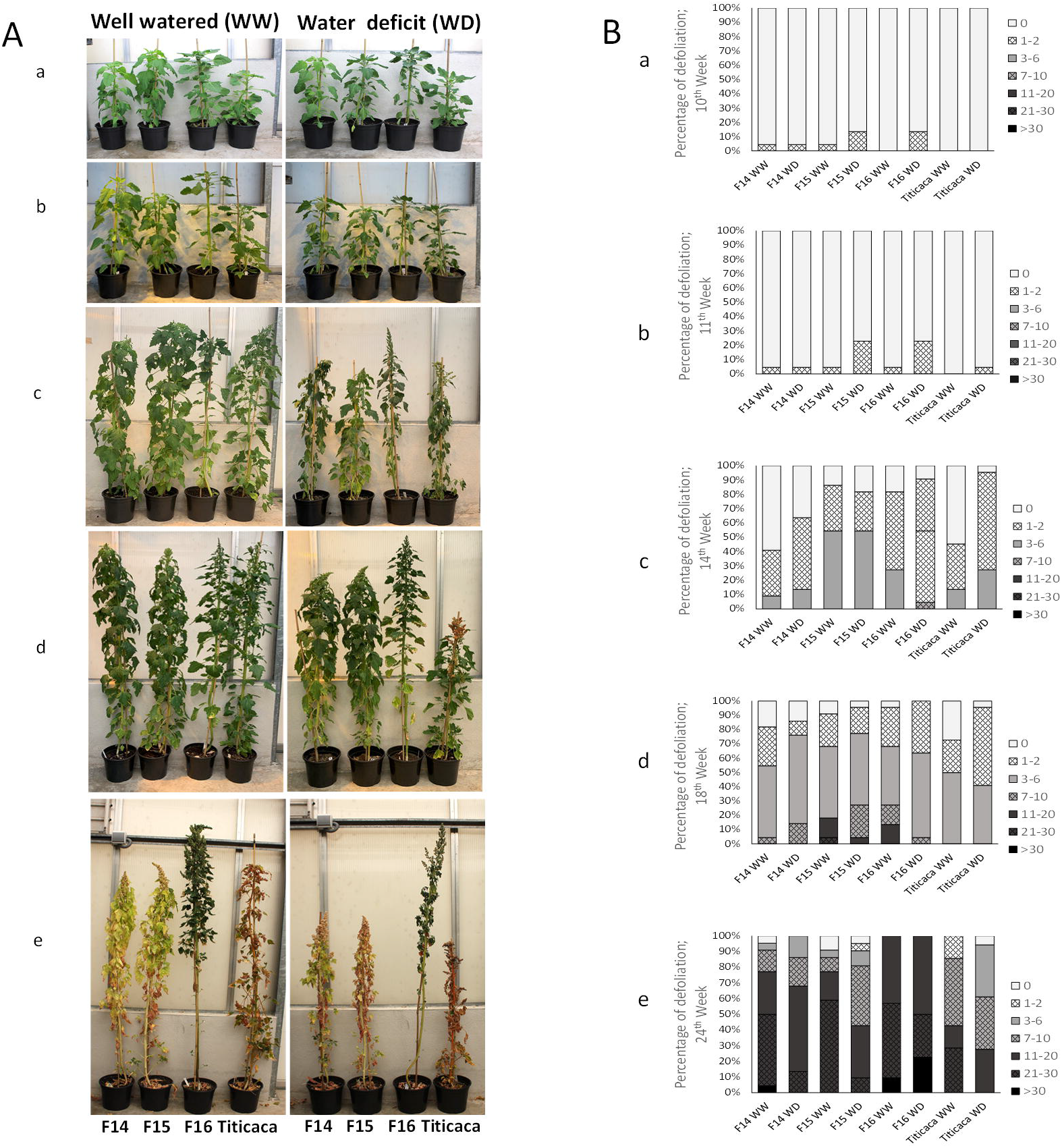
Time course images and defoliation rates of different quinoa cultivars grown under different water treatments (WW or WD) throughout development. (**A**) Images of the quinoa plants including both water treatments (in the left image WW conditions, or in the right image WD conditions) and the four cultivars analyzed in this study (F14, F15, F16 and Titicaca) together with (**B**) the defoliation rates by the phenological stage. Letters indicate the week number after seed sowing as follows: a: 10^th^ week; b: 11^th^ week; c: 14^th^ week; d: 18^th^ week and e: 24^th^ week.

### 3.2 Plant physiological responses

The chlorophyll index was measured weekly on upper fully expanded leaves (Fig. 5A) and also was taken at different parts in the plant (upper, middle, and lower part) (Supplementary Fig. 3). The chlorophyll index in the upper fully expanded leaves of the plants was calculated and a 3-Way ANOVA was performed to evaluate the influence of the three factors. The developmental stage (*p*<0.001), the cultivar (*p*<0.001), the water treatment (*p*=0.001), the interaction between the developmental stage and the cultivar (*p*=0.003), and the interaction between the cultivar and the water treatment (*p*=0.006), influenced this parameter. Furthermore, significantly higher levels of chlorophyll were found in WD plants compared to WW plants. Also, an increment of chlorophyll was observed till the 16^th^ week, followed by a progressive decrease until the end of the experiment (seed mature stage). When focusing on the differences between cultivars, it was observed that F16 was able to maintain the chlorophyll levels constant during development, independently of the water treatment, and showed a higher chlorophyll index than the rest of cultivars independently of the water treatment. When comparing water treatments within each cultivar, it was noted that F15 showed higher chlorophyll levels under WD than in WW, differences that were kept up to the 22^nd^ week. At the seed filling stage, Titicaca WW plants showed higher levels of chlorophyll compared with WD plants (Fig. 5A).

**Figure 5.**
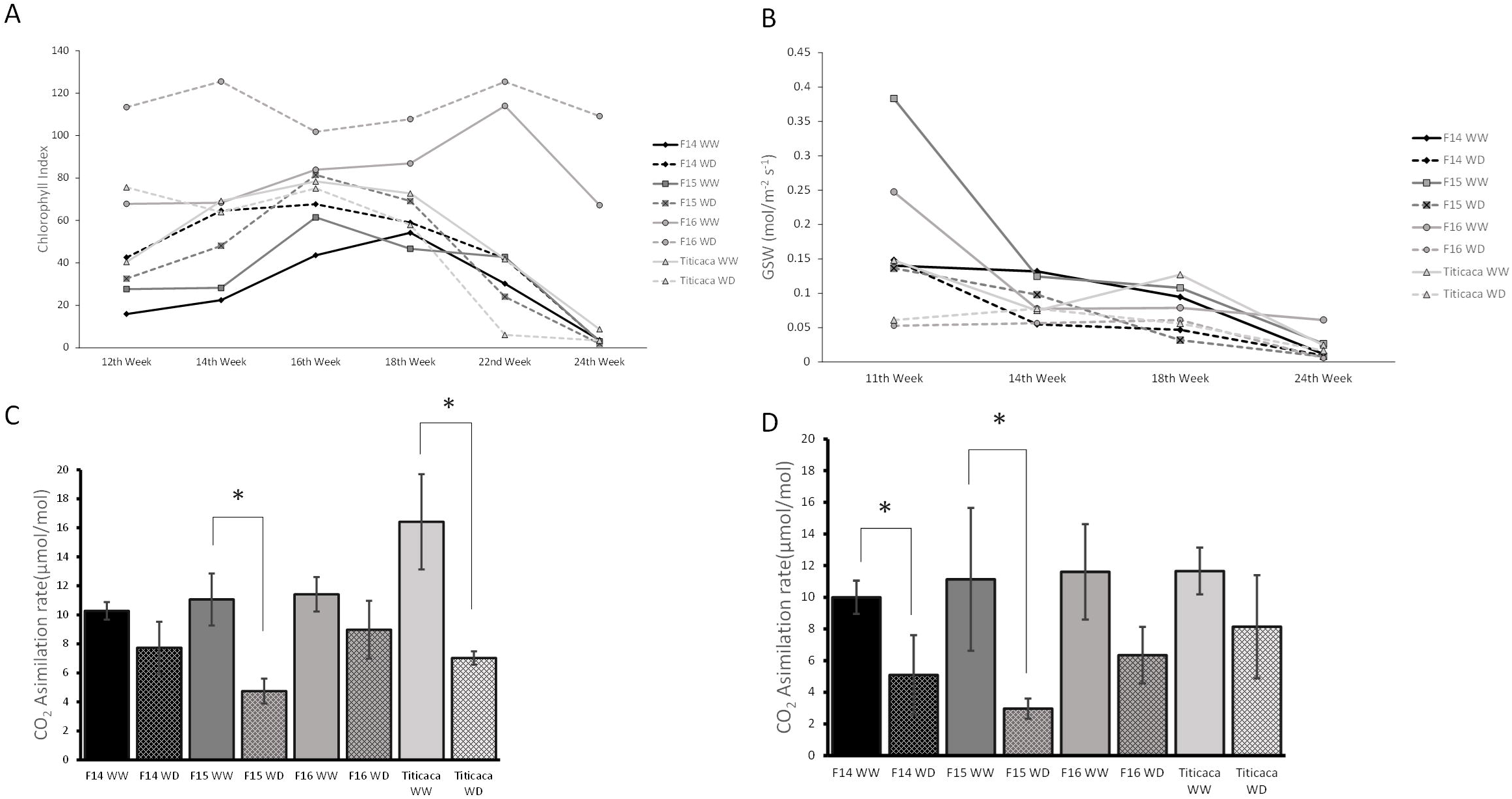
Photosynthetic-related parameters in quinoa cultivars growing under well-watered (WW) or water stress conditions (WD). (**A**) The chlorophyll index, (**B**) the stomatal conductance (GSW (mol/m^-2^ s^-1^)), (**C**) the CO_2_ assimilation rates at vegetative state (μmol/mol) (Week 13^th^) and (**D**) the CO_2_ assimilation rates at seed filling state (μmol/mol) (Week 17^th^) measured in the upper fully expanded leaves of the plant. The statistical analyses performed for the chlorophyll index and the stomatal conductance data analysis are presented in Supplementary Table 3. Error bars in panels C and D represent the standard deviation of the mean value. Asterisks (*) indicate statistically significant differences among cultivars subjected to different water treatments (WW or WD), following a pairwise comparison (t-Student when the data followed a normal distribution or U Mann-Whitney when the data did not follow a normal distribution) at a *p*-value <0.05.

When comparing the chlorophyll index among the different parts of the plant, it was observed that the only cultivar that showed a gradient in the chlorophyll index, from the upper part of the plant to the lower part, was F16, while the rest of cultivars (independently of the water treatment), showed similar chlorophyll levels in the upper and middle leaves, being these higher than the lower leaves’ chlorophyll index (Supplementary Fig. 3).

*S*tomatal conductance (GSW) was measured throughout the experiment (Fig. 5B). All the factors analysed, including the cultivar (*p*<0.001), treatment (*p*<0.001), and developmental stage (*p*<0.001), and their interactions (*p*<0.001), influenced this parameter. A decrease in GSW was shown as the crop was growing, being this parameter higher in WW plants than WD plants, and, in general terms, among cultivars, being higher in the F15 cultivar. On the other hand, GSW was decreasing gradually over time for all cultivars, independently of the water treatment. Nonetheless, differences were observed among cultivars. For instance, Titicaca GSW behaved similarly between water treatments and maintained GSW levels constant until their decrease on the 24^th^ week (at the physiological maturity stage). Also, F16 GSW showed similar values from flowering (14^th^ week) till the end of the experiment when differences appeared between WW and WD conditions.

CO_2_ assimilation rates were analysed at two critical development stages (at vegetative and seed filling stages) (Fig. 5C and 5D). By performing a 3-Way ANOVA analysis, the significant factors influencing this parameter were the water treatment (*p* <0.001), the cultivar (*p* <0.001), the developmental stage (*p*=0.006), the interaction between cultivar and water treatment (*p*=0.023) and the interaction among the developmental stage, the water treatment and the cultivar (*p*=0.001). Generally, higher assimilation rates were observed under WW treatment than in WD. Besides, higher assimilation rates were found at the vegetative stage compared to the seed filling stage, and differences among cultivars revealed that Titicaca and F16 plants were the ones presenting higher CO_2_ assimilation rates compared to F15 plants. Pair comparisons showed distinct patterns depending on the developmental stage. Thus, in F15 plants, CO_2_ assimilation rates were lower at the vegetative stage under WD conditions, while in WW Titicaca plants, the levels of CO_2_ assimilation rates were lower at the seed filling stage. Moreover, when comparing water treatments, differences appeared in the cultivar F15 at the vegetative and seed filling stages, with higher rates under WW conditions (Fig. 5C and 5D), Titicaca at the vegetative stage, with larger rates found under WW conditions (Fig. 5C), and F14 at the seed filling stage, with larger rates found under WW conditions (Fig. 5D).

Water use efficiency (WUE) was calculated considering the photosynthetic rates (A) and the stomatal conductance (gs), as the ratio A/gs (Supplementary Fig. 4). WUE of F15 and Titicaca cultivars did not change with the water treatment at both developmental stages (vegetative and seed filling stage). On the contrary, F16 and F14 cultivars subjected to WD showed higher levels of WUE than the WW plants (at both development stages) (Supplementary Fig. 4).

When the levels of WUE were related to the amount of water applied to keep the water regimes equal on the soil (Supplementary Fig. 1B) it was observed that the cultivars F15 and Titicaca were the ones presenting larger water consumption rates during development, contrary to the response observed in F14 and F16 plants. Particularly F15, despite being the cultivar receiving larger amounts of water, the soil water content (SWC) of F15 pots remained lower compared to the SWC of the rest of cultivars (Supplementary Fig. 1A).

Chlorophyll fluorescence measurements were taken to evaluate the status of the photosynthetic membrane (Kalaji et al., 2016) (Fig. 6). Among the parameters evaluated, the efficiency of the photosystem II (*ΦPSII*) remained constant throughout the experiment, with a small decrease observed in the 14^th^ week and a sharp decrease at seed maturation (24^th^ week) (Fig. 6A). No differences were observed in *ΦPSII* between water treatments (*p*=0.430) (WW and WD) nor cultivars (*p*=0.199). Nonetheless, this parameter was influenced by the developmental stage (*p*<0.001), the interaction between the developmental stage and the water treatment (*p*=0.002), the developmental stage, and the cultivar (*p*<0.001), and by the interaction between the water treatment and the cultivar (*p*=0.002). Besides, differences appeared when evaluating changes of this parameter linked to the developmental stage, being F16 the only cultivar that did not show significant differences throughout development. Also, when comparing by water treatments, F16 and F14 cultivars showed higher values of *ΦPSΓΓ* at WD than at WW at the inflorescence stage, prior to flowering (11^th^ and 14^th^ week respectively). At later stages, WW Titicaca and F16 cultivars presented higher levels of *ΦPSΓΓ* than under WD conditions.

**Figure 6.**
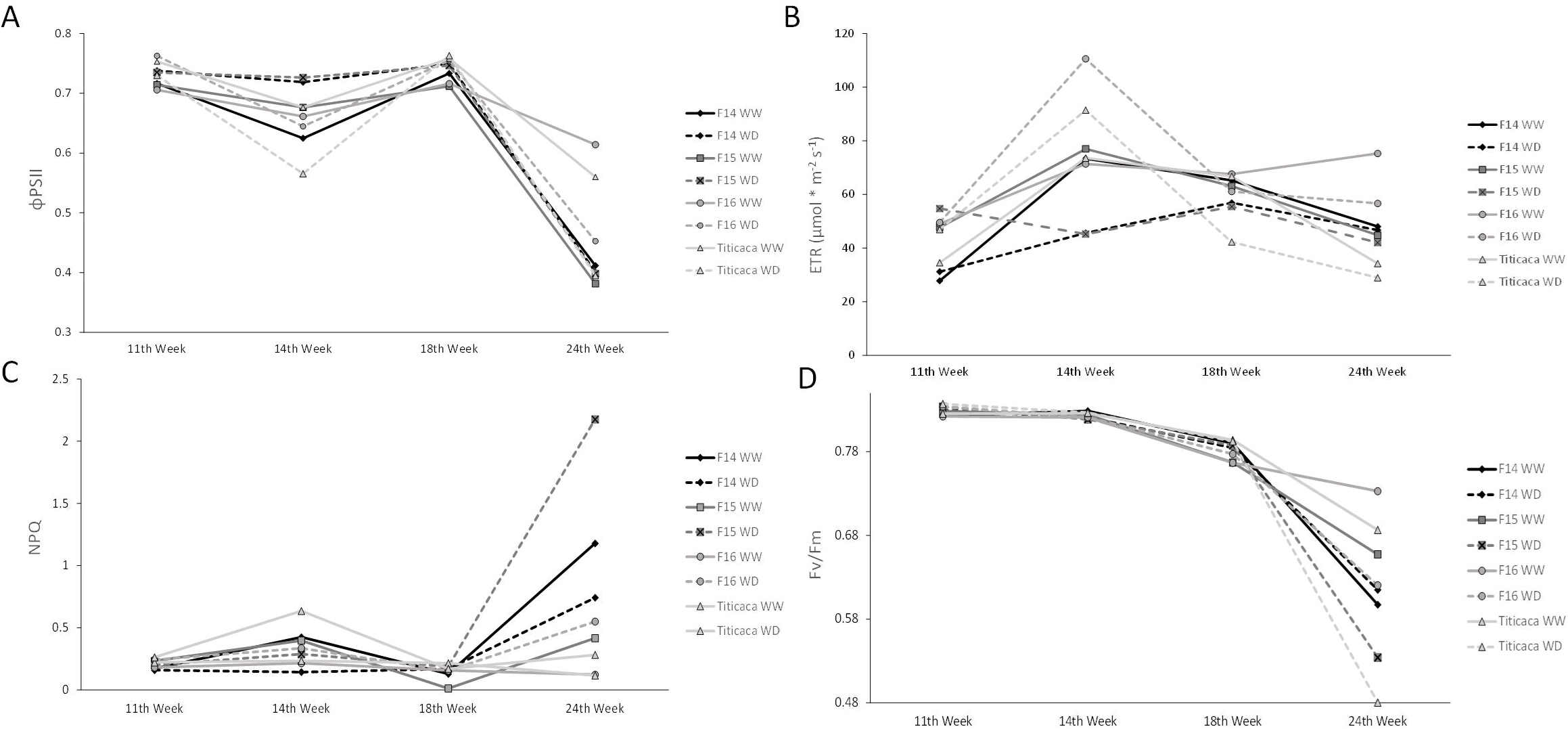
Chlorophyll fluorescence-related parameters measured throughout development in different quinoa cultivars subjected to different water treatments (WW or WD). The chlorophyll fluorescence-related parameters included: (**A**) the efficiency of the PSII (*Φ*PSII) (**B**), the electron transport rate (ETR (μmol * m^-2^ s^-1^)) (**C**), the non-photochemical quenching (NPQ) (**D**), and the maximum efficiency of photosystem II (Fv/Fm) and were all measured throughout development in the different cultivars evaluated under different water regimes. Each water condition is represented by either continuous (WW) or dashed lines (WS) and each genotype with rhombus (F14), squares (F15), circles (F16) or triangles (Titicaca). The statistical analyses performed for these parameters are presented in Supplementary Table 3.

Another chlorophyll fluorescence associated parameter, the electron transport rate (*ETR*), was influenced by the developmental stage (*p*<0.001), the cultivar (*p*<0.001), the interaction between the developmental stage and the cultivar (*p*<0.001), and by the interaction among the developmental stage, the cultivar, and the water treatment (*p*<0.001) (Fig. 6B). *ETR* did not show differences between water treatments but did show differences depending on the cultivar. In line with this, F16 showed higher *ETR* values compared to the other cultivars. In general, differences associated with the developmental stage were observed, with an increase at the 14^th^ and 18^th^ weeks and a later decrease at the 24^th^ week, reaching again 11^th^ week *ETR* values. When comparing *ETR* values in each developmental stage, this parameter showed specific differences. For example, F15 and Titicaca WD plants presented lower levels at weeks 14 and 18, while F16 WD plants kept *ETR* levels constant during development. The pair comparison between WW and WD plants revealed developmental-dependent differences. Thus, at pre-anthesis, no differences were found between WW and WD in F16 (*p=*0.863) and Titicaca (*p*=0.436) cultivars, but higher *ETR* values were found at WW for the cultivars F14 (*p*=0.011) and F15 (*p*=0.001). On the contrary, at the seed mature stage, no differences were found between treatments (Fig. 6B).

The non-photochemical quenching (*NPQ*) did not reveal a significant influence of the factors analysed (*p*=0.060) (Fig. 6C). In general terms, NPQ remained constant during the experiment although it showed a small decrease at the 18^th^ week in all cultivars and for both water treatments. No differences were observed between WW and WD or among cultivars, although the pair comparison showed particular differences, such as the higher NPQ at WD in F14 compared to F14 WW plants (at pre-anthesis) or the higher NPQ values showed by F16 WD compared to WW at seed mature stage (Fig 6C).

The maximum quantum yield of PSII (*Fv/Fm*) was also quantified (Fig. 6D). A 3-Way ANOVA test showed an influence of the water treatment (*p*<0.001) and the developmental stage (*p*<0.001), including the interactions between these factors and the cultivar (*p*<0.001 for all the interactions except for the interaction between the cultivar and the water treatment which was *p*=0.011). Differences appeared in *Fv/Fm* levels between water treatments, with higher values under WW compared to WD. No differences appeared among cultivars. Considering the developmental stage, a reduction of *Fv/Fm* along the phenological development was observed. Analysing the differences between water treatments and among cultivars over time, it was observed that *Fv/Fm* decreased over time starting at the 14^th^ week under WW and the 11^th^ week under WD. When comparing by water treatment in each cultivar, all the *Fv/Fm* values were similar except for the cultivar F14, in which both water treatments showed a small increase in *Fv/Fm* at the 14^th^ week. Furthermore, pairwise comparisons during development were performed. In moments prior to anthesis, differences between WW and WD treatments were observed for the cultivar F14 (*p*=0.044). In WW F14, *Fv/Fm* values were higher than in WD plants. At the final stages of seed maturation, no differences were observed between water treatments in the F14 cultivar (*p*=0.656), but higher levels of *Fv/Fm* were observed in WW plants compared to WD plants in F15, F16, and Titicaca cultivars (*p*=0.006, *p*=0.005, and *p*=0.001, respectively) (Fig. 6D).

### 3.3 Seed Yield and Harvest Index (HI)

Seed yield was determined per cultivar and treatment. All WW plants showed higher seed yields than WD plants (F14 +27,8%, F15 +43,2%, and Titicaca +43%, on average) except for F16, which did not show differences between treatments (Fig. 7A). No differences in seed yield were found among cultivars when growing under WD. Under WW conditions the only difference among cultivars appeared between Titicaca and F16 plants, with Titicaca presenting higher yields than F16 cultivar.

**Figure 7.**
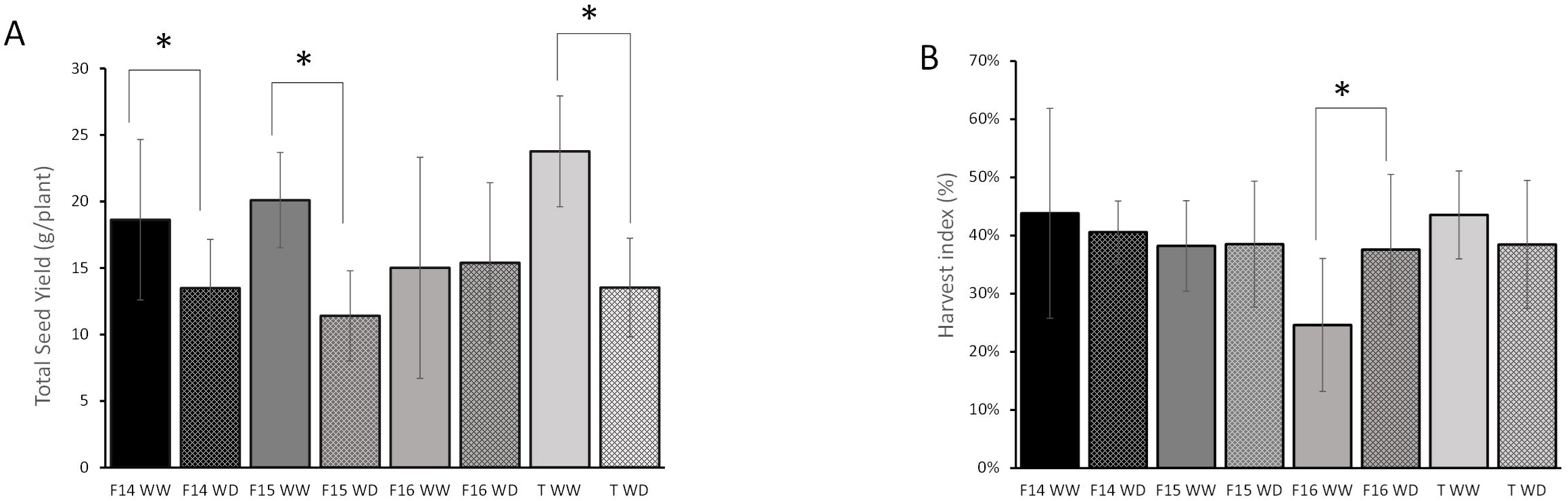
Total seed yield and harvest index of the quinoa cultivars subjected to both water treatments (WW or WD). (**A**) Total Seed Yield (g of seeds/plant) and (**B**) Harvest Index (%) were determined at the end of the experiment for each cultivar (F14, F15, F16, and Titicaca) subjected to both water treatments (WW, black and grey columns) or WD (dashed black and grey columns). Error bars represent the standard deviation of the mean value. Asterisks (*) show statistically significant differences among cultivars subjected to different water treatments, following a pairwise comparison (t-Student when the data followed a normal distribution or U Mann-Whitney when the data did not follow a normal distribution) at a *p*-value <0.05.

Significant differences in the HI only occurred in the cultivar F16, in which WW F16 plants showed lower HI values than F16 WD (Fig. 7B). At harvesting, the only cultivar that did not show biomass penalties due to WD was F16 (Supplementary Fig. 5). The rest of the cultivars reduced their plant biomass under water stress by decreasing the leaf, stem, and/or seed biomass.

## 4. Discussion

Plants trigger different mechanisms to overcome abiotic stress depending on the species. Quinoa is well known for being an abiotic stress-tolerant crop, including drought (Jacobsen et al., 2003). Gómez-Pando et al. (Gómez-Pando et al., 2019) attributed quinoa’s drought tolerance to three main mechanisms: drought escape, which is related to the shortening of the life cycle (Jacobsen et al., 2003); drought avoidance, which can be achieved by optimising water absorption and water loss through a vigorous root system, defoliation, and stomatal regulation (Jensen et al., 2000); and drought physiological tolerance, acquired through tissue elasticity and osmolyte regulation (Bascuñán-Godoy et al., 2016; Cutler et al., 1977). Nonetheless, a decrease in photoprotection mechanisms has been described in this plant when subjected to water stress (Bosque Sanchez et al., 2006).

Regarding drought escape strategies, a reduction of yield associated with water deficits has been reported in many different staple crops, such as wheat, maize, or rice (Daryanto et al., 2016; Kumar et al., 2014). This has been linked to a lifespan shortening consequence of the changes in the plant phenology. For instance, in maize, it was shown that water stress resulted in the shortening of the vegetative stage accelerating, consequently, flowering and reducing the grain-filling period, which ended in a grain yield decrease (Samarah, 2005; Shavrukov et al., 2017). Drought stress applied during flowering or the grain-filling period can also shorten the reproductive stage of barley and rice causing grain yield penalties (Kadam et al., 2018; Pantuwan et al., 2002; Samarah, 2005). In line with this, a negative effect of WD on seed yield in quinoa has been previously reported (Geerts et al., 2008). Geerts et al. (Geerts et al., 2008) applied severe WD at different developmental stages revealing that the milky seed stage (during seed filling) was the most sensitive phase to drought followed by flowering. Besides, it was observed that drought may cause the shortening of quinoa life cycle in the field (Jacobsen et al., 2003). However, to date, there are very few studies performed in quinoa analysing the specific physiological and phenological responses to drought, particularly under long-term water stress, depicting the genotypic control in this respect. In this regard, our study confirms the genotype-dependency associated with WD response in this crop. For instance, the phenology of genotype F14 showed an opposing response to F15, F16, or Titicaca cultivars, increasing its lifespan under WD compared to WW conditions (Fig.1), which highlights the importance of the genetic factor as determinant of the water-stress response in quinoa.

In the current study, the genotype that generally showed higher WD tolerance (considering drought avoidance strategies like a lower water consumption, the maintenance of CO_2_ assimilation rates, and the stability of the photosynthetic membrane together with lesser seed yield penalties) was F16. Furthermore, in other crops such as wheat or lettuce, small decreases in water availability can result in higher photosynthetic rates maintaining yields due to the improvement in WUE under drought as observed in some quinoa cultivars in this study (Molina-Montenegro et al., 2011; Van Den Boogaard et al., 1997) (Supplementary Fig. 4). Higher WUE can be achieved by reducing the stomatal conductance, while maintaining the photosynthetic capacity (Jacobsen et al., 2009). An increase in WUE under WD conditions has also been described in quinoa, and it was related to the stomata closure that resulted in the maintenance of leaf water potential, keeping active photosynthesis (Geerts et al., 2008). In here, a similar response was observed in some cultivars (Supplementary Fig. 4). In fact, the F16 cultivar was able to increase WUE under WD, reducing the stomatal conductance (in comparison with WW conditions) and keeping CO_2_ assimilation rates close to WW conditions, ultimately maintaining seed yield and increasing HI under water-limiting conditions (Figs. 5 and 7B). The enhanced WUE and tolerance to WD of this genotype were also associated with leaf area reduction under drought, also observed in F15 (Supplementary Fig. 2). This strategy has been developed by other important crops such as wheat under WD (Barraclough et al., 1989).

Another trait that could have contributed to enhancing water-stress tolerance by reducing transpiration in the cultivar F16 was its defoliation rate, which was higher when compared to the rest of the cultivars and was more pronounced at the final stages of development (Fig. 4B). Higher defoliation rates in this cultivar were observed in the lower parts of the plant leading to a concentration of leaves (source tissue) in the upper part of the plant, that were also horizontally disposed around the inflorescence (sink tissue), as it can be observed in Fig. 4, while the other cultivars did not show such response. Furthermore, the chlorophyll gradient was much more marked in F16 which could reflect a senescence induction in lower leaves prior to defoliation (Supplementary Fig. 3). Besides, plant height was another growth-related trait evaluated in this study, decreasing under stress in all the genotypes analysed (Fig. 2). An inhibitory effect that has been also observed in other crop species (Çakir, 2004), and, in the case of F16, was less pronounced and was related to a higher HI while not being related to seed yield penalties (Fig.7).

The plant architecture was very variable among genotypes, and this can impact water tolerance (Tognetti et al., 2010). Following the quinoa growth habits defined by Rojas and Pinto (Rojas & Pinto, 2013), the cultivar F16 would fit within the growth habit 1, in which the number of branches is reduced, F14 could be classified between habits 1 and 2, F15 between habit 2 and 3, and Titicaca within the habit 4, these last presenting larger ramification number, leaves, and therefore, being more susceptible to water loss due to higher total leaf surface. This highlights a possible association in quinoa between water-stress tolerance and the growth habit.

Considering all the parameters evaluated in this study, a schematic summary is presented in Fig. 8 in which a distinction can be made between tolerant and sensitive genotypes in quinoa based on the water use, different morphological and photosynthetic-related parameters, and agronomical traits. Thus, a water stress tolerant genotype would increase WUE under stress conditions, reducing water consumption by lowering its leaf area, increasing its defoliation rates and chlorophyll index, and concentrating the leaves in the upper part of the plant closer to the sink tissue (inflorescence). In line with this, a genotype fitting in the growth habit 1 described by Rojas and Pinto (Rojas & Pinto, 2013) would show an innate advantage facing WD, since a plant architecture presenting fewer and smaller branches would allow for lower leaf total surface. Besides, in the tolerant genotypes, a reduced GSW would avoid water loss, without showing a large inhibition of photosynthesis, thus maintaining seed yield parallel to a HI increment. Accordingly, considering all these aspects, we could say that quinoa has the potential to present drought-mediating mechanisms which are genotype-dependent (Jacobsen, 2003; Jacobsen et al., 2003, 2009, 2013).

**Figure 8.**
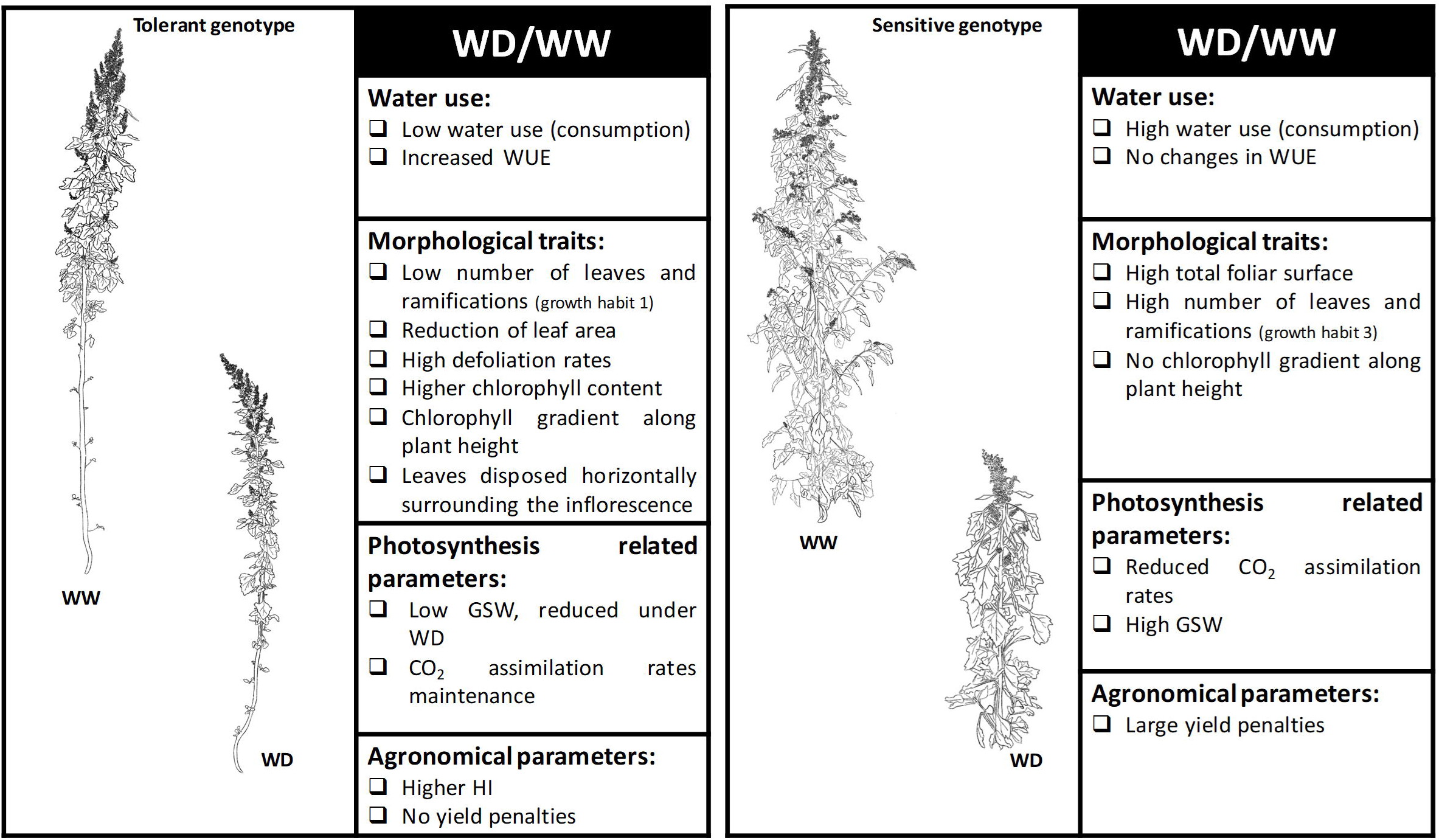
Main morphological, physiological and agronomical traits associated to two contrasting phenotypes: water tolerant versus water sensitive quinoa genotypes. This schematic model highlights the main characteristics associated to a water tolerant or a water sensitive quinoa genotype when facing drought conditions. These characteristics were classified in four main groups: water use, morphological traits, photosynthesis related parameters and agronomical parameters. The model was based on the results presented by the different genotypes analyzed in the current study that reflected a differential response to water stress, being F16 a genotype that would fit in the water tolerant genotype group, and F15 one that would mostly fit in the water sensitive genotype group.

Environmental degradation of arid regions is often associated with the loss of vegetation cover, soil, and water resources, which could also result from agricultural practices (Clarke & Noin, 1998). In this sense, to minimize these impacts, agriculture could bet on high-yielding resilient nutritious crops such as quinoa (Jaikishun et al., 2019). Furthermore, within the current global environmental context, linked to limited water availability in some extensive agricultural areas, finding cultivars that require less water while keeping productivity is mandatory. In this regard, our study reveals that certain cultivars, like F16, possess characteristics here proposed as promising for rain-fed areas (Fig. 8) since, despite not being the most productive genotype under WW conditions (compared with the other cultivars analysed), it was the cultivar requiring less amount of water (Supplementary Fig. 1), preserving better this limited resource while maintaining its yield. On the contrary, the F15 cultivar, which presented similar yields under WW conditions than WW F16, suffered severe seed yield penalties under WD (Fig.5), and, presented larger water consumption rates (Supplementary Fig. 1B), thus aligning closer to the sensitive model phenotype (Fig. 8). Interestingly, some authors have argued that a bred crop developed with improved WUE, cannot attain high yield potential, similarly to what was observed in the present study (Blum, 2005) (Fig. 7A). The rationale here is based on the premise that if breeding retains characteristics associated with yield potential increments (i.e. larger leaf areas) when drought stress occurs, the same traits selected to improve productivity are detrimental for water preservation. In line with this, it might be a matter of reaching a balance between productivity and water requirement, which should be a preferred criterion when selecting quinoa varieties for each agronomical context, especially those destined to rain-fed conditions. Nonetheless, it should not be forgotten that, in the field, several biotic and abiotic factors may act simultaneously inducing stress (Ben Rejeb et al., 2014; Ramegowda & Senthil-Kumar, 2015; Reguera et al., 2012). Therefore, even though F16 would be an optimal cultivar for rain-fed conditions according to our findings, its longer life cycle could negatively impact its performance in Mediterranean climates if flowering or seed filling stages occurred later in the season, coinciding with high temperatures (Matías et al., 2021).

Overall, we can conclude that the cultivars here evaluated presented different mechanisms to cope with long-term water stress, including changes in phenology, morphology, or in their physiological response. All these genotype-dependent responses to WD conditions resulted in yield penalties in most of the cultivars tested except for F16 (Fig. 7A), which might be the most promising genotype to grow under water-limiting conditions. Thus, considering the current climate prospects in which certain agricultural areas will suffer more frequent drought episodes (European Commission, 2019; FAO, 2016, 2022) together with the need of re-valuing rain-fed agriculture, particularly important in the Mediterranean area (Araus, 2004; Jacobsen et al., 2013) the selection of more WUE quinoa cultivars is crucial. In line with this, it is required that we better comprehend the plant physiological responses associated to water stress using experimental designs able to mimic field conditions, to ensure the reproducibility of the results. The application of long-term water stress when analysing plant physiological responses might be tedious but the experimental conditions are closer to what we can find in nature. Altogether, we believe that this work will significantly contribute to broadening our understanding regarding how quinoa responds to long-term water stress highlighting genotype-related differences that will allow the selection of the best adapted genotypes for water-limiting environments.

## Abbreviations

ETR: electron transportation rate
DW: dry weight
FW: fresh weight
GSW: stomatal conductance
HI: harvest index
Fv/Fm: maximum quantum yield of photosystem II
ΦPSII: efficiency of photosystem II
NPQ: non-photochemical quenching
SWC: soil water content
WD: water-deficit
WW: well-watered.

## Author Contributions

M.R., J.M., I.M.G., and S.G.R. conceived and planned the experiments. I.M.G., S.G.R., M.O., and M.R. carried out the experiments. M.R., I.M.G., S.G.R., J.M., M.O., V.C., and L.B. contributed to the interpretation of the results. M.R., I.M.G., and S.G.R. took the lead in writing the manuscript. All authors provided critical feedback and helped shape the research, analysis, and manuscript.

## Data Availability

All data supporting the findings of this study are available within the paper and within its supplementary materials published online

## Acknowledgments

The authors greatly thank Susana Vilariño (Algosur, Spain) for her support helping to conceive and plan the experiments and for providing, together with Dr. Sven Jacobsen (Quinoa Quality, Denmark), the quinoa seeds used in this study. The authors would also like to specially thank Rosendo López García (CBGP (UPM, INIA-CSIC) for his technical assistance and help in the greenhouse.

## Supplementary data

Supplementary data are available at JXB online.

Table S1. Leaf number and ramification number of quinoa plants at vegetative stage.

Table S2. Statistical analysis performed for the plant height parameter.

Table S3. Statistical analysis performed for the photosynthetic-related parameters.

Table S4. Statistical analysis performed for chlorophyll fluorescence-related parameters.

Fig. S1. Soil water content and water supply in the quinoa pots used throughout the experiment.

Fig. S2. Leaf area of newly fully expanded leaves of quinoa growing under two water treatments (WW or WD).

Fig. S3. Chlorophyll index gradient of leaves determined at two developmental stages (vegetative stage and at seed filling stage) in different quinoa genotypes growing at two different water conditions (WW and WD).

Fig. S4. Water use efficiency (WUE) of different quinoa cultivars growing under two wáter regimes (WW or WD).

Fig. S5. Total dry plant biomass at harvesting of different quinoa cultivars growing under two different water treatments (WW or WD).

